# Static and Dynamic Aspects of Cerebro-Cerebellar Functional Connectivity are Associated with Self-reported Measures of Impulsivity: A Resting-State fMRI Study

**DOI:** 10.1101/2020.01.20.912295

**Authors:** Majd Abdallah, Nicolas Farrugia, Valentine Chirokoff, Sandra Chanraud

**Affiliations:** CNRS UMR 5287, Aquitaine Institute of Cognitive and Integrative Neuroscience, University of Bordeaux, France; Laboratory of Neuroimaging and Daily Life, EPHE, PSL Research University, Bordeaux, France; Electronics Department, Lab-STICC - IMT Atlantique, Brest, France

**Keywords:** Cerebellum, impulsivity, cerebro-cerebellar networks, functional connectivity, strength, temporal variability

## Abstract

Converging evidence from human and animal studies predict a possible role of the cerebellum in impulsivity. However, this hypothesis has not been thoroughly investigated within the framework of functional connectivity (FC). To address this issue, we employed resting-state fMRI data and two self-reports of impulsivity (UPPS-P and BIS/BAS) from a large group of healthy young individuals (N=134). We identified cerebral and cerebellar resting-state networks, and evaluated the association of static (strength) and dynamic (temporal variability) aspects of cerebro-cerebellar FC with different elements of self-reported impulsivity. Our results revealed that the behavioral inhibition and approach systems (BIS/BAS) were inversely associated with basal ganglia-cerebellar and fronto-cerebellar FC strength, respectively. In addition, we found that lack of premeditation was inversely associated with the temporal variability of FC between the cerebellum and top-down control networks that included sub-regions of the prefrontal cortex, precuneus, and posterior cingulate cortex. Moreover, we found that sensation seeking was associated with the temporal variability of FC between the cerebellum and networks that included cortical control regions and sub-cortical reward regions: the basal ganglia and the thalamus. Together, these findings indicate that the cerebellum may contribute to different forms of impulsivity through its connections to large-scale control and reward networks.

## 1 Introduction

Impulsivity is a multi-dimensional personality construct that is present to different degrees in healthy individuals as well as diverse psychiatric populations (Bakhshani, 2014). It is defined as the tendency to initiate actions dominated by spontaneity and urgency with little to no reflection or consideration of consequences (Madden and Bickel, 2010; Bakhshani, 2014). Studies have shown that highly impulsive individuals engage more frequently in risky behaviors such as substance use, gambling, aggressive behavior, and increased food intake (Dickman, 1990; Moeller et al., 2001; Madden and Bickel, 2010). Impulsivity is usually assessed using self-report questionnaires addressing two broad processes: inhibitory control and sensitivity to reward and punishment (Dawe et al., 2004). Inter-individual differences in these processes can arise from different genetic and neural origins that are not fully understood (Khadka et al., 2014). However, monoaminergic cortico-striatal systems have been shown to mediate the different constructs of impulsivity, and alterations to these systems have been observed in highly impulsive individuals and clinical groups with impulsive symptomatology (Dalley et al., 2011; Mitchell and Potenza, 2014; Fineberg et al., 2014). Recently, however, a hypothesis for the involvement of the cerebellum in impulsivity has been advanced based on numerous findings from human and animal studies. These findings indicate that the cerebellum might be involved in terminating or initiating actions by regulating cerebral networks that mostly include prefrontal and striatal regions (Miquel et al., 2019).

Originally thought of as a sensorimotor structure, the cerebellum is now known to be functionally diverse and involved in higher-order cognitive processes (Leiner et al., 1986; Buckner et al., 2011; Buckner, 2013). Studies have confirmed that the majority of the cerebellum maps onto association regions in a manner that mirrors the cerebral asymmetries for cognition, language, and attention (Habas et al., 2009; Buckner, 2013). Converging evidence have indicated that the cerebellum provides automatic predictions via forward models to most cortical processes, in a uniform manner, including decision making, inhibitory control and reward processing (Deverett et al., 2018; Miquel et al., 2019). Studies have identified closed loops connecting the cerebellum to cortical and sub-cortical regions that sub-serve control and reward such as the ventral tagmental area (VTA), prefrontal cortex (PFC), orbito-frontal cortex (OFC), anterior cingulate cortex (ACC), basal-ganglia, and insula (Caligiore et al., 2017; Moreno-Rius and Miquel, 2017; Carta et al., 2019). In addition, studies have observed alterations in cerebro-cerebellar functional connectivity (FC) in disorders exhibiting impulsive symptomatology such as alcohol use disorder (Chanraud et al., 2011), borderline personality disorder (De Vidovich et al., 2016), and certain neurodevelopmental disorders (Miller et al., 2010). Although these findings, among others, predict a possible role of the cerebellum in impulsive behavior, a direct assessment of this hypothesis is still lacking in the FC literature. One way to test this hypothesis is by exploring potential associations between cross-sectional differences in different elements of impulsivity and different aspects of cerebro-cerebellar resting-state functional connectivity (rs-FC).

Resting-state FC analysis mainly involves estimating summary measures of the overall FC strength between different neuronal populations across a scanning period of a few minutes, and assessing potential associations with behavioral factors (den Heuvel and Pol, 2010; Sporns, 2018). This has proven to be highly informative in uncovering the neural substrates underlying certain behaviors, cognitive abilities, and personality traits in healthy and clinical populations. However, it is now argued that this “static” approach masks information embedded in the dynamic nature of the brain (Calhoun et al., 2014; Lurie et al., 2018). Accordingly, recent FC studies have begun exploring the temporal dynamics of brain activity and have pointed to the presence of flexible non-random FC configuration (brain states) that transiently appear during task and rest conditions (Allen et al., 2012; Vidaurre et al., 2017). FC dynamics have been shown to be highly related to higher-order cognitive processes such as learning and memory (Fatima et al., 2016; Braun et al., 2015). In addition, it is argued that dynamic FC measures complement and, in some cases, outperform static FC measures in explaining behavioral factors (Liéegeois et al., 2019). However, static and dynamic FC measures explain different aspects of behavior, and information from both can explain more variance than either alone (Ramos-Nunñez et al., 2017; Liéegeois et al., 2019). Major advances have been made in understating how brain networks dynamically interact and impact behavior. However, little is known about the dynamics of the cerebellum and cerebro-cerebellar networks and their relation to behavior. That being said, we believe that by exploiting the dynamics of cerebro-cerebellar FC, we can gain deeper insight into the functional repertoire of the cerebellum. Therefore, the aim of this study is to evaluate static and dynamic aspects of cerebro-cerebellar FC in order to test the hypothesis of a cerebellar involvement in impulsivity.

We investigate this hypothesis using a large dataset (N = 134), comprising resting state functional magnetic resonance imaging (rs-fMRI) data and measures of impulsive behavior from the Max Planck institute Mind-Brain-Body database (Mendes et al., 2019). In this sample, impulsivity is assessed using the UPPS-P impulsive behavior scale and the behavioral inhibition and approach systems (BIS/BAS) that capture different forms of impulsive behavior. First, we decompose the rs-fMRI data into separate cerebral and cerebellar resting-state networks (RSNs) and extract the corresponding BOLD time-series. We then estimate summary measures of the overall FC strength and temporal variability between cerebral RSNs of interest and the cerebellum using robust methods. Since cerebro-cerebellar networks form closed-loops, cerebellar outputs are believed to directly modulate singular network components rather than affecting large-scale complex processes. However, indirect connections are also believed to expand the influence of the cerebellum (Sokolov, 2018). Therefore, we use full and partial correlations as measures of FC in both the static and dynamic FC analyses. Finally, we test for potential associations between measures of cerebro-cerebellar FC and impulsivity using general linear models (GLMs) and permutation testing, and assess the validity of significant findings using 5-folds cross-validation.

## 2 Materials and Methods

### 2.1 Sample and behavioral Measures

The study included participants from the Neuroanatomy and Connectivity protocol (N&C) which is part of the larger Max Planck institute Mind-Brain-Body database. All data were acquired and made publicly available by Max Planck Institute of Human Cognitive and Brain Sciences in Leipzig,Germany (Mendes et al., 2019). All included participants were healthy with no past nor present signs of any neuropsychiatric condition. The dataset included two widely-used scales measuring different impulsivity-related traits: the UPPS-P impulsivity scale and the Behavioral Inhibition/Approach Systems Scales (BIS/BAS). The UPPS-P is a self-report designed to measure impulsive behavior across the five factor model of personality: sensation seeking, negative urgency, positive urgency, lack of premeditation, and lack of perseverance. These sub-scales are not considered direct measures of trait impulsivity, but rather measures of distinct personality traits that can lead to impulsive behavior (Whiteside et al., 2005). The BIS/BAS included summary measures of two general motivational systems argued by theorists to underlie behavior: a behavioral inhibition (avoidance) system measuring sensitivity to punishment or unpleasant cues, and a behavioral approach system measuring sensitivity to desirable cues (i.e. goals and rewards) (Gray, 1991). The included BAS score was defined as the sum of three sub-scales: BAS drive, BAS fun seeking, and BAS reward responsiveness. The normality of the scores was tested using the Shapiro-Wilik test of normality and rank-based inverse Gaussian transform was performed to counteract departures from normality. Taken together, these measures represented a sufficient set of variables covering different aspects of trait impulsivity in which the cerebellum has a hypothesized role; inhibitory control and reward processing. More information on the dataset, the complete set of behavioral measures, and the inclusion/exclusion criteria are found in (Mendes et al., 2019).

### 2.2 Resting-state fMRI data and pre-processing

The openly available rs-fMRI dataset originally included preprocessed data from 188 subjects. We excluded, due to a gap in the age distribution, 26 subjects that were older than 55. Additional 28 subjects were excluded for missing or corrupt resting-state or behavioral data from source. The final sample included 134 healthy young subjects (46% female) aged 20-40 years. The resting-state fMRI data acquisition parameters are found in full detail in (Mendes et al., 2019). In summary, four resting-state fMRI scans were acquired for each individual in axial orientation using T2*-weighted gradient-echo echo planar imaging (GE-EPI) with multi-band acceleration. Sequences were identical across the four runs, with the exception of varying slice orientation and phase-encoding direction. The phase-encoding direction was anterior–posterior (AP) for runs 1 and 3, and posterior-anterior (PA) for runs 2 and 4. The complete set of parameters was set as follows: voxel size = 2.3 mm isotropic, FOV = 202 × 202 mm^2^, imaging matrix = 88 × 88, 64 slices with 2.3 mm thickness, TR = 1400 ms, TE = 39.4 ms, flip angle = 69^*◦*^, echo spacing = 0.67 ms, bandwidth = 1776 Hz/Px, partial fourier 7/8, no pre scan normalization, multiband acceleration factor = 4657 volumes, duration = 15 min 30 s per run. Individuals were instructed to remain awake, during the resting-state scan, with their eyes open and to fixate on a cross-hair.

The preprocessing pipeline is described in details in (Mendes et al., 2019) and the corresponding python code can be obtained from https://github.com/NeuroanatomyAndConnectivity/pipelines/tree/master/src/lsd_lemon. In summary, the preprocessing steps included 1) removal of the first 5 volumes from each of the 4 resting state runs, 2) rigid body alignment to the first volume using FSL MCFLIRT to obtain transformation parameters; 3) field-map unwarping using FSL-FLIRT and FSL-FUGUE to estimate transformation parameters for distortion correction; 4) co-registration to each subject’s structural scan via FreeSurfer’s boundary-based registration to estimate transformation parameters for co-registration; 5) application of the transformation parameters to each volume in the four resting state runs in one interpolation step; 6) six motion parameters, their first-order derivatives, and outliers from Nipype’s rapidart algorithm were included as nuisance regressors in a GLM; 7) the *aCompCor* method to regress out WM and CSF related signals; 8) Band-pass filtering [0.01Hz −0.1Hz]; and 9) normalization to MNI152 2mm space. All included subjects show relatively low in-scanner motion (mean frame-wise displacement (Power et al., 2012)) (mean-FD*<*0.5 mm) across all 4 scans. Fully pre-processed data were obtained and downloaded from https://ftp.gwdg.de/pub/misc/MPI-Leipzig_Mind-Brain-Body/derivatives/.

### 2.3 Group-ICA spatial maps and time-series extraction

#### 2.3.1 Cerebellum-only group-ICA

In order to estimate the resting-state networks (RSNs) in the cerebellum, we performed cerebellum-only group ICA after computing a group average cerebellar mask in MNI space. This approach has proven to be more sensitive and robust in capturing cerebellar RSNs as compared to whole-brain group-ICA (Dobromyslin et al., 2012; Kipping et al., 2016). The group-ICA was performed using the GIFT toolbox http://mialab.mrn.org/software/gift/ using different implemented tools. First, cerebellar rs-fMRI data from all subjects were demeaned and used in a subject-level principal components analysis (PCA) to reduce the dimensionality to 100 subject-level principal components (retaining *>* 99% of the variance in the data). A second PCA was performed at the group-level to further reduce the dimensionality to 25 principal components. The choice of number of components was based on findings from previous studies that identified between 7 and 20 cerebellar RSNs using different data-driven methods (Buckner et al., 2011; Bernard et al., 2012; Kipping et al., 2016; Wang et al., 2016). Next, the Infomax algorithm was used to estimate the cerebellar independent components (ICs). The stability of these estimates was assessed by running the ICASSO algorithm 20 times, as described and implemented in the GIFT toolbox, and the most stable configuration was automatically chosen as the final result. Finally, the components’ time-series were extracted using group information guided ICA, or GIG-ICA, due to its ability to estimate subject-specific ICs and time-series with better accuracy and inter-individual correspondence than other methods such as GICA, GICA3, and dual-regression (Du et al., 2017; Salman et al., 2019). GIG-ICA uses the group-level spatial maps as guidance to estimate individual subject-specific ICs and time-series. Detailed description of this algorithm can be found in (Du and Fan, 2013). Components that exhibited strong activation near the grey matter/white matter/cerebro-spinal fluid borders and/or exhibited noisy patterns with no functional relevance were discarded. Only those components that exhibited unilateral/bilateral activation patterns in the grey matter and had relevance to well-known cerebellar functional clusters were selected as cerebellar RSNs of interest (Buckner et al., 2011). Finally, the cerebellar RSNs time-series were standardized to have a mean of 0 and standard deviation of 1 for each subject in each resting-state run.

#### 2.3.2 Cerebral group-ICA

A similar approach to the cerebellum-only group-ICA was performed in order to estimate cerebral RSNs. First, a group average mask in MNI space, that did not include the cerebellum, was computed. Then cerebral rs-fMRI data were demeaned and used in a two-stages PCA to estimate 120 subject-level principal components (retaining *>* 99% of the variance of the data) and 30 group-level principal components. The choice of a low-order model was driven by our interest in large-scale brain RSNs that were also suitable for subsequent FC analysis methods in terms of dimensionality, complexity, and interpretability. We used the infomax algorithm and ICASSO to estimate and automatically select the most stable set of group independent components, followed by GIG-ICA to estimate subject-specific components’ time-series. We eliminated from the analysis components showing strong activation near the edges and in the white matter. Only those components that exhibited activation patterns in the grey matter and that had relevance to well-known large-scale functional networks were included as RSNs of interest (Yeo et al., 2011). Finally, the cerebral RSNs time-series were standardized to have a mean of 0 and standard deviation of 1 for each subject in each resting-state run.

### 2.4 Functional Connectivity Analysis

#### 2.4.1 Static functional connectivity

After identifying cerebral and cerebellar RSNs and extracting their time-series, we performed a static FC analysis in order to estimate subject-level FC matrices. Specifically, we estimated full and partial correlation matrices using the Ledoit-Wolf estimator implemented in the nilearn and scikit-learn python packages (Ledoit and Wolf, 2004; Abraham et al., 2014). The obtained FC matrices were Fisher r-to-z transformed to stabilize the variance, and corrected for the effective number of degrees of freedom according to Bartlett’s method which controls for the effect of auto-correlation on the estimation of FC (Bartlett, 1946; Afyouni et al., 2019). Next, in each resting-state run, we computed the FC strength of cerebral RSNs of interest as the sum of their positively weighted cerebellar edges and then averaged the values across four resting-state runs for each subject. Negatively weighted edges were not considered due to the lack of consensus and ambiguity concerning their nature, interpretation, and means of analysis (Fornito et al., 2016; Sporns and Betzel, 2016; Hallquist and Hillary, 2018). The cerebral RSNs of interest included cortical and sub-cortical networks that might be involved in control and reward processes, namely orbito-frontal, salience, attention, fronto-parietal, default-mode, basal-ganglia, and thalamus networks. This yielded a FC strength vector of size 134 subjects *×* 16 RSNs in each of the full and partial correlation cases.

#### 2.4.2 Dynamic functional connectivity

In order to model whole-brain FC dynamics we applied a hidden Markov model (HMM) on the temporally concatenated BOLD timeseries from all subjects using the hidden Markov models multivariate auto-regression (HMM-MAR) toolbox https://github.com/OHBA-analysis/HMM-MAR (Vidaurre et al., 2017). The HMM is a window-less dynamic FC approach that bypasses the limitations of sliding-windows by being directly applied to the BOLD time-series. In addition, the method uses variational Bayesian inference and minimisation of free-energy to update and estimate group-level brain states, each described as a multivariate Gaussian distribution with a mean representing a spatial activation pattern, and a covariance matrix representing a FC pattern (Vidaurre et al., 2017). The Bayesian inference process implemented within the HMM also provides the probability of occurrence of each state at each time point, along with the Viterbi path which represents the most likely sequence of states (Quinn et al., 2018). These were used to compute the subject-specific frequency of occurrence of each state defined as the number of times this state was visited across the whole scanning period. The HMM requires a pre-specified number of states, so we assumed a fixed number of six states as a compromise between a lower-model order (5 states) and higher-model orders (8, 10 and 12 states) after performing a stability analysis using all configurations (see supplementary figure S3). The number of states within the HMM paradigm provides a description of the dataset at a certain granularity where a higher number means finer temporal resolution (Quinn et al., 2018). It is worth noting that we were mostly interested in FC changes rather than changes in the absolute signal, hence brain states were only defined by their full and partial correlation matrices. Partial correlation matrices were obtained by inverting the covariance matrices which were automatically regularized within the Bayesian framework of the HMM (Ryali et al., 2016).

##### Subject-specific brain states and temporal variability

In order to estimate a descriptive summary metric of cerebro-cerebellar FC dynamics, we explored the manifestation of the states at the subject-level. Particularly, we performed an additional iteration of the Bayesian inference process at the subject-level, given the initial group-level estimates and the subjects’ BOLD data as prior information when updating and re-inferring the states. The subject-level dynamic FC matrices were Fisher r-to-z transformed and used to compute the state-wise FC strength of cerebral RSNs of interest as the sum of their positively weighted cerebellar edges, in a similar fashion to the static FC analysis. We then computed the temporal variability of FC between each cerebral RSN of interest and the cerebellum as the unbiased frequency-weighted standard deviation of state-wise FC strength values across the states for each subject. This was accomplished using the built-in MATLAB ***std*** function that treated state-wise FC strength values as weighted samples and the subject-specific states frequency of occurrence values as weights (see figure 1, panel C). Similar to the static FC analysis, cerebral RSNs of interest included cortical and sub-cortical networks that might be involved in control and reward processes, which yielded a FC temporal variability vector of size 134 subjects × 16 RSNs in each of the full and partial correlation cases.

**Figure 1:**
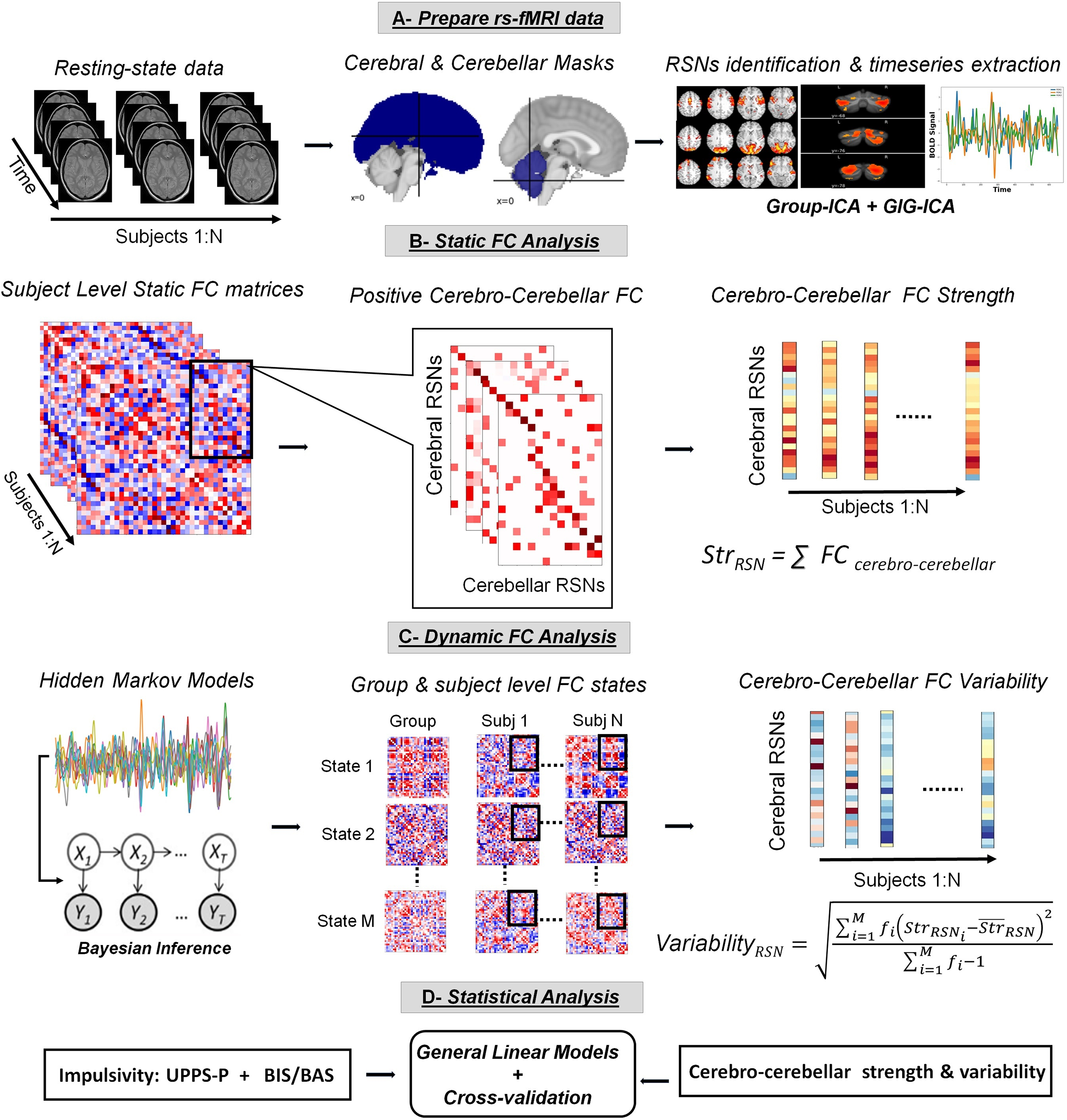
The overall methodology architecture. A) Preprocessed BOLD data from all subjects were used to estimate group-level cerebral and cerebellar RSNs after separate masks in MNI space. B) Static FC analysis included the estimation of subject-level full and partial correlation matrices, the extraction of positive cerebro-cerebellar edges, and the calculation of FC strength *Str*_*RSN*_ as the sum of the positively weighted cerebro-cerebellar links of each cerebral RSN. C) Dynamic FC analysis included the estimation of group and subject-level dynamic brain states via HMM, extraction of positive cerebro-cerebellar edges (not shown explicitly), and the calculation of FC temporal variability *V ar*_*RSN*_ as the unbiased frequency (*f*_*i*_) weighted standard deviation of state-wise cerebro-cerebellar FC strength values across multiple states for each subject. D) Statistical analysis using GLMs and cross-validation

The rationale behind this approach was based on the observation that static FC matrices across subjects highly resembled the frequency-weighted mean of the dynamic FC matrices: cosine similarity > 0.98 for full correlation matrices and *>* 0.94 for partial correlation matrices (see supplementary figure S4). Therefore, static FC could be considered as a superposition of dynamic FC states identified via HMM, as was also observed by Karapanagiotidis et al. (2018). In other words, the dynamic FC patterns could be considered as non-random transient deflections from the static FC pattern at shorter time-scales. More detailed theoretical and practical information on the use of HMMs can be found in Baker et al. (2014); Ryali et al. (2016); Vidaurre et al. (2017); Quinn et al. (2018), and https://github.com/OHBA-analysis/HMM-MAR/wiki/User-Guide.

##### Assessing the robustness of FC dynamics

In order to confirm the presence of robust and genuine dynamic FC patterns in the rs-fMRI data, we generated 100 null datasets from a multivariate Gaussian distribution fitted to the real data of each individual subject as in (Vidaurre et al., 2017). According these authors, the static correlations in the simulated data are similar to those in the real data but are presumed to be stationary. We performed HMM, with unchanged parameters, on each set of the null data and extracted a metric that allowed us to compare the results to those obtained in the real data. One important metric that has been used in previously is the maximum fractional occupancy. Generally, fractional occupancy is defined as the proportion or percentage of time each state is visited by each subject, whereas maximum fractional occupancy is simply the maximum proportion of time spent by each subject visiting the most occurring state (Vidaurre et al., 2017). High maximum fractional occupancy values closer to 1 indicate that a single state describes the entirety of the data and hence the absence of dynamics in FC. In this context, we compared the distributions of maximum fractional occupancy values in the real and null datasets to assess the presence/absence of genuine FC dynamics in the rs-fMRI data.

### 2.5 Associations between cerebro-cerebellar FC and impulsivity scores

We used multivariate general linear models (GLMs) to test the effect of the impulsivity measures on cerebro-cerebellar FC measures while controlling for age, gender, and mean frame-wise displacement. We performed non-parametric permutation testing with 10,000 permutations and a maximum z-statistic procedure to obtain family-wise adjusted p-values for each test. This method provides strong control of Type I error rates without being highly conservative as conventional correction techniques (e.g. Bonferroni) (Winkler et al., 2014). We reported significant associations with *p <* 0.05 FWE-adjusted. Next, in order to assess the validity of significant findings, we used repeated stratified 5-folds cross-validation to split the entire sample into training (80% of data, 107 subjects) and testing (20% of data, 27 subjects) sub-samples where the percentages of males and females were preserved in each split. The cross-validation scheme was repeated 100 times with a different randomization in each repetition. In this case, all subjects took part in the testing phase across folds and repetitions. GLMs were fitted to the training data and then used to predict the outcome in the unseen data. We included only those variables that exhibited significant associations in the previous step (i.e. GLMs fitted to the entire data) while controlling for age, gender, and mean frame-wise displacement. Finally, we reported the median values of the explained variance obtained in the training (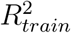) and testing (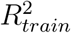) sub-samples across all folds/repetitions.

## 3 Results

### 3.1 Behavioral and Demographic Data

Summary statistics of demographic and behavioral data are provided in table 1, whereas the partial correlations between the different impulsivity measures are provided in table S1 in supplementary material. Subjects were originally grouped by age bins each spanning 5 years, but we considered the lower bound of each bin as the age of each subject. Variables were standardized to have a mean equal to zero and standard deviation equal to one. We assessed multicollinearity in the data using the variance inflation factor (VIF) approach. Most variables were found to have a *VIF <* 2 except for the UPPS-P negative and positive urgency sub-scales. Accordingly, in order to avoid potential multicollinearity due to the strong association between the two sub-scales (Student’s *t* = 11.34, *r* = 0.68, *p <* 10^−15^), we used factor analysis to obtain one urgency factor while conserving a sufficient amount of variance. The final set of self-reported impulsivity measures included six sub-scales: urgency, lack of premeditation, lack of perseverance, sensation seeking, behavioral inhibition system (BIS), and behavioral approach system (BAS).

**Table 1:**
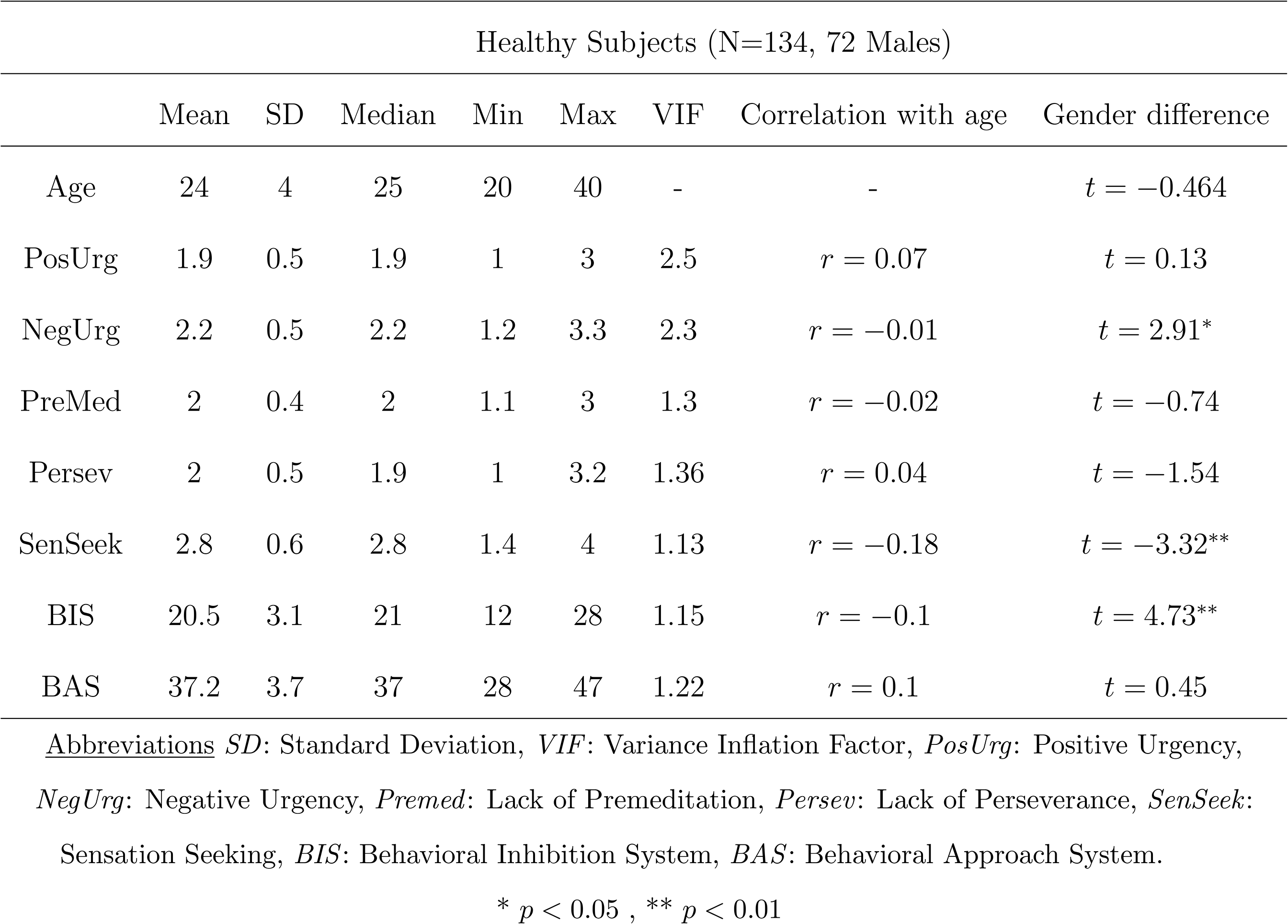
Demographic and behavioral summary statistics

| Healthy Subjects (N=134, 72 Males) |  |  |  |  |  |  |  |  |
| --- | --- | --- | --- | --- | --- | --- | --- | --- |
|  | Mean | SD | Median | Min | Max | VIF | Correlation with age | Gender difference |
| Age | 24 | 4 | 25 | 20 | 40 | - | - | $t = -0.464$ |
| PosUrg | 1.9 | 0.5 | 1.9 | 1 | 3 | 2.5 | $r = 0.07$ | $t = 0.13$ |
| NegUrg | 2.2 | 0.5 | 2.2 | 1.2 | 3.3 | 2.3 | $r = -0.01$ | $t = 2.91^*$ |
| PreMed | 2 | 0.4 | 2 | 1.1 | 3 | 1.3 | $r = -0.02$ | $t = -0.74$ |
| Persev | 2 | 0.5 | 1.9 | 1 | 3.2 | 1.36 | $r = 0.04$ | $t = -1.54$ |
| SenSeek | 2.8 | 0.6 | 2.8 | 1.4 | 4 | 1.13 | $r = -0.18$ | $t = -3.32^{**}$ |
| BIS | 20.5 | 3.1 | 21 | 12 | 28 | 1.15 | $r = -0.1$ | $t = 4.73^{**}$ |
| BAS | 37.2 | 3.7 | 37 | 28 | 47 | 1.22 | $r = 0.1$ | $t = 0.45$ |

Abbreviations *SD*: Standard Deviation, *VIF*: Variance Inflation Factor, *PosUrg*: Positive Urgency, *NegUrg*: Negative Urgency, *Premed*: Lack of Premeditation, *Persev*: Lack of Perseverance, *SenSeek*: Sensation Seeking, *BIS*: Behavioral Inhibition System, *BAS*: Behavioral Approach System.
\* $p < 0.05$ , \*\* $p < 0.01$

### 3.2 Group-ICA RSNs Estimation

#### 3.2.1 Cerebellar RSNs

Resting-state data from the cerebellum were decomposed into 25 ICs, out of which 14 ICs were identified as cerebellar RSNs and shown in Figure 2. The identified RSNs were arranged into groups of putative functional domains based on their anatomical or functional properties and their overlap with well-established cerebellar clusters (Buckner et al., 2011). The functional clusters were: sensory-motor, visual, attention, salience, fronto-parietal, default-Mode, and language. However, two cerebellar RSNs, whose spatial activation maps were well situated in the GM, did not overlap with well-known cerebellar clusters. These were labelled as “Vermis” and “Crus-I/II” based on the anatomical landmarks that overlap with their spatial maps. In addition, taking into consideration the contralateral representation of large-scale networks in the cerebellum, labels of unilateral cerebellar RSNs were inverted. For instance, if activation was mostly localized in the left anterior cerebellum, the naming would be cerebellar right motor network (Cer-rMot) due to the inverted somato-motor map present in the anterior cerebellar lobe.

**Figure 2:**
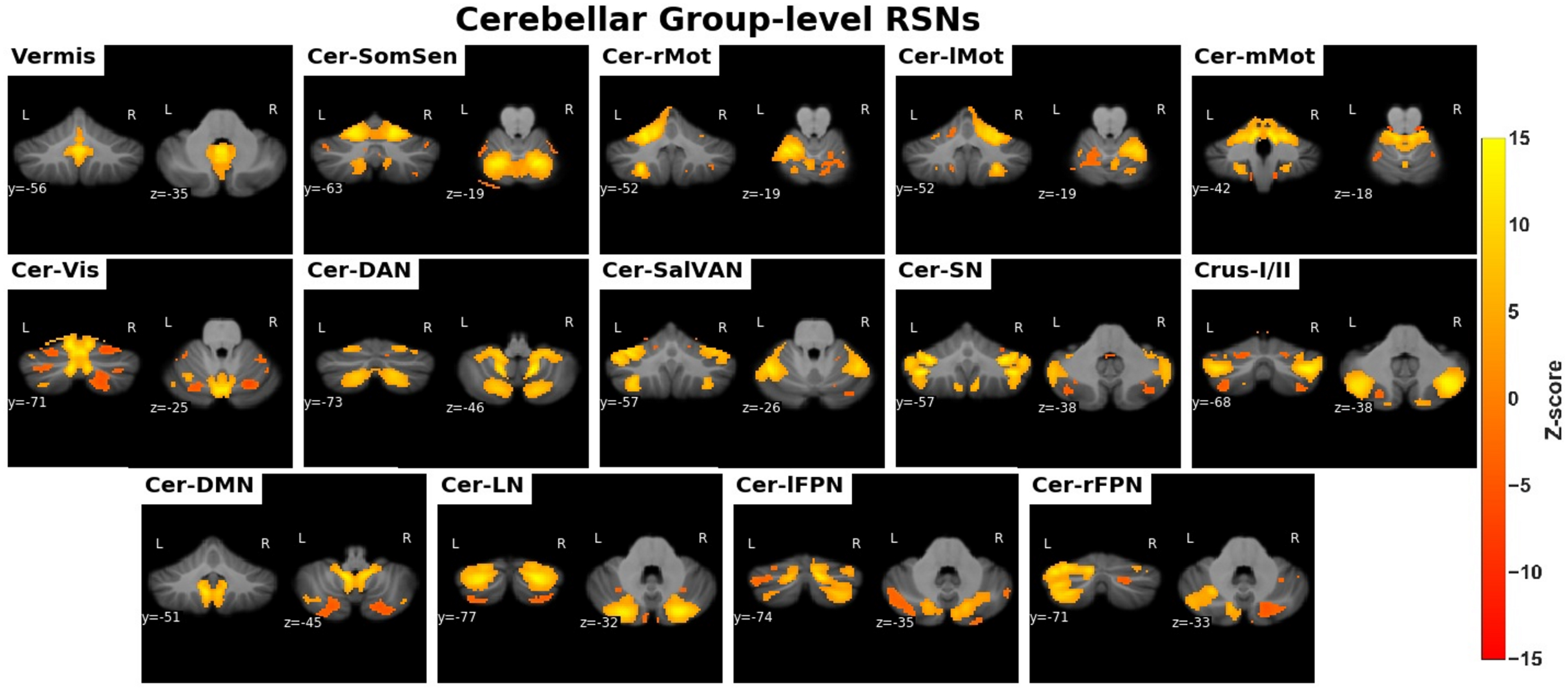
The identified cerebellar RSNs. <u>Abbreviations</u> *Cer-SomSen*: Cerebellar somato-sensory network, *Cer-rMot*: Cerebellar right motor Network,*Cer-lMot*: Cerebellar left-motor network, *Cer-mMot*: Cerebellar medial motor network, *Cer-Vis*: Cerebellar visual network, *Cer-DAN*: Cerebellar dorsal attention network, *Cer-SalVan*: Cerebellar salience/ventral attention network, *Cer-SalEx*: Cerebellar salience network, *Cer-DMN*: Cereballar default-mode network, *Cer-LN*: Cerebellar language network, *Cer-lFPN*: Cerebellar left fronto-parietal network, *Cer-rFPN*: Cerebellar right fronto-parietal network

#### 3.2.2 Cerebral RSNs

The data from the cerebral cortex and sub-cortex were decomposed into 30 ICs out of which 25 ICs were identified as RSNs (see Figure 3) based on visual inspection of the localization of activation in the grey matter (GM). The retained RSNs were arranged into groups of putative functional domains based on their anatomical and functional properties: sub-cortical, somato-motor, visual, auditory, attention, salience, fronto-parietal, and default Mode. Finally, 39 cerebral and cerebellar RSNs were retained for further FC analysis.

**Figure 3:**
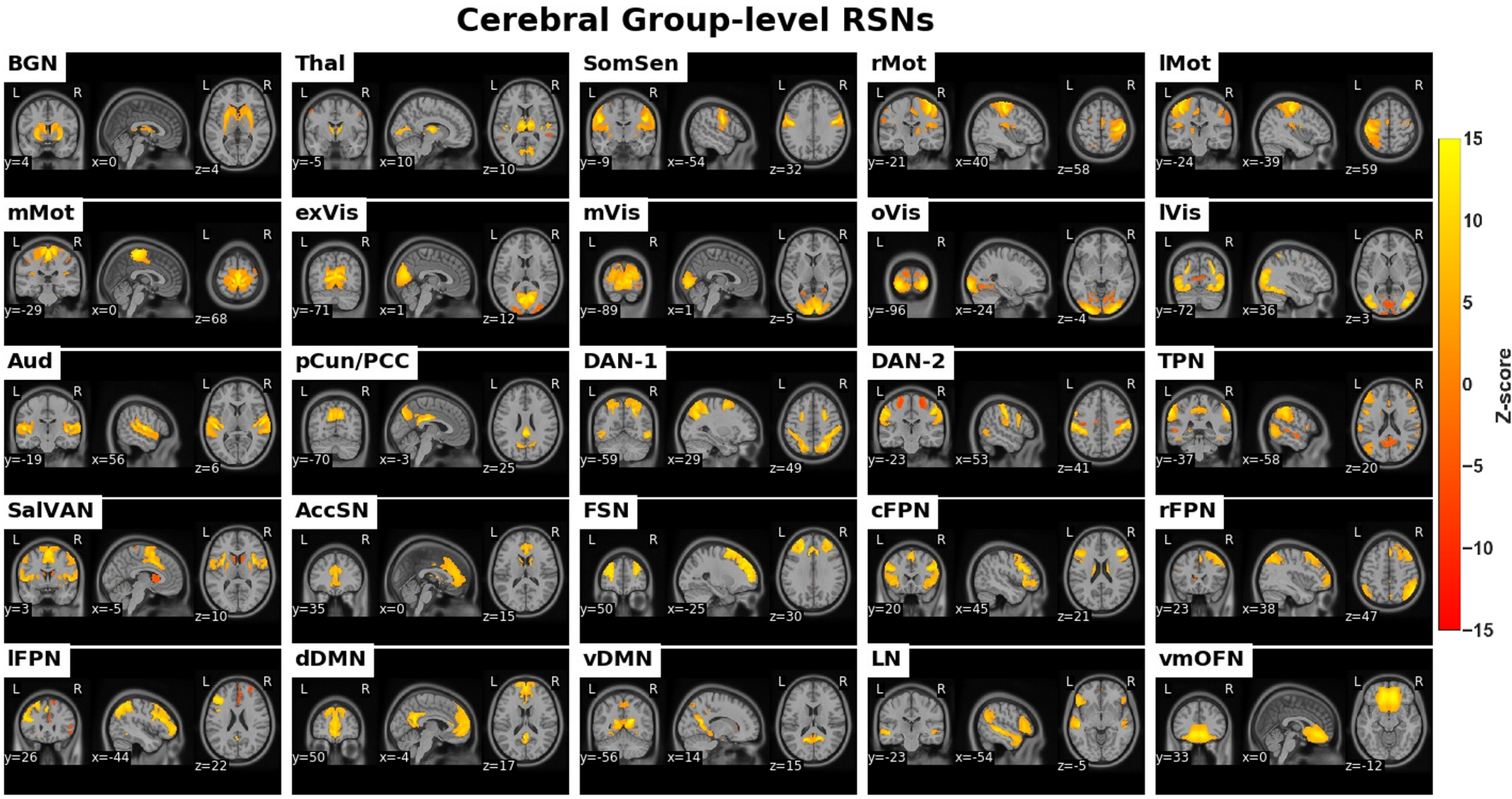
The identified large-scale cerebral RSNs. <u>Abbreviations</u> *BGN*: Basal-ganglia network, *Thal*: Thalamus, *SomSen*: Somatosensory network, *rMot*: Right motor Network, *lMot*: Left-motor network, *mMot*: Medial motor network, *exVis*: Extra-striat visual network, *mVis*: Medial Visual network, *oVis*: Occipital Visual Network, *lVis*: Lateral visual network, *Aud*: Auditory network, *pCun/PCC*: Precuneus/Posterior cingulate cortex network, *DAN*: Dorsal Attention Network, *TPN*: Task positive network, *SalVAN*: Salience-ventral attention network, *AccSal*: Anterior cingulate cortex salience network, *FSN*: Frontal salience network, *cFPRN*: Central fronto-parietal network, *rFPN*: Right fronto-parietal network, *lFPN*: Left fronto-parietal network, *dDMN*: Dorsal default-mode network, *vDMN*: Ventral default-mode network, *LN*: Language network, *vmOFN*: Ventro-medial orbito-frontal network

### 3.3 Cerebro-Cerebellar FC strength and Impulsivity

The statistical details of significant associations between impulsivity scores and cerbero-cerebellar FC strength are reported in table 2, whereas scatter plots are illustrated in supplementary figure S1; panel A. We used figure 4 to demonstrate the group average static FC matrix as an overview of the strongest cerebro-cerebellar direct connections. It clearly shows the well documented topographic dichotomy of anterior motor vs posterior non-motor cerebellum. We reported significant results by showing z-scores, standardised regression coefficients (*β*), FWE-adjusted p-values, and the median of explained variance values (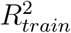) and (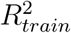) across folds/repetitions. Results revealed a significant inverse association between the behavioral approach system (BAS) scale and the FC strength between the frontal salience network (FSN) and the cerebellum (*z* = −3.1, *β* = −0.29, *p* = 0.033, 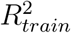 = 0.08, 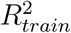 = 0.073) obtained from partial correlation matrices. In addition, we identified a significant inverse association between the behavioral inhibition system scale (BIS) scale and the FC strength between the basal-ganglia network (BGN) and the cerebellum (*z* = −3, *β* = −0.31, *p* = 0.038, 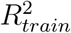 = 0.07, 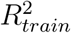 = 0.068) obtained using partial correlations. No significant associations were observed when quantifying static FC using full correlations.

**Table 2:** GLMs and cross-validation results

| Impulsivity scale | Network | FC Metric | FC type | $z$ | $\beta$ | p-value (FWER) | $R^2_{train}$ | $R^2_{test}$ |
| --- | --- | --- | --- | --- | --- | --- | --- | --- |
| BIS | BGN-cerebellum | strength | partial | -3 | -0.31 | 0.038 | 0.07 | 0.068 |
| BAS | FSN-cerebellum | strength | partial | -3.1 | -0.29 | 0.033 | 0.08 | 0.073 |
| SenSeek | BGN-cerebellum | variability | full | 3.1 | 0.3 | 0.037 | 0.068 | 0.063 |
|  | Thal-cerebellum | variability | full | 3.3 | 0.32 | 0.019 | 0.078 | 0.074 |
|  | FSN-cerebellum | variability | full | 3.3 | 0.32 | 0.019 | 0.078 | 0.08 |
|  | pCun/PCC-cerebellum | variability | full | 3.6 | 0.35 | <.01 | 0.093 | 0.086 |
| PreMed | FSN-cerebellum | variability | full | -3.5 | -0.34 | <.01 | 0.092 | 0.09 |
|  | pCun/PCC-cerebellum | variability | full | -3.7 | -0.36 | <.01 | 0.11 | 0.096 |

Abbreviations *FSN*: Frontal salience network, *pCun/PCC*: Precuneus/Posterior-cingulate cortex network
*BGN*: Basal-ganglia network, *Thal*: Thalamus network, *BAS*: Behavioral approach system, *BIS*:
Behavioral inhibition system, *SenSeek*: UPPS-P Sensation Seeking, *PreMed*: UPPS-P Lack of premeditation, $z$ : Computed z-statistics, $\beta$ : Standardised regression coefficient, *p-value*: Family-wise error adjusted p-value, $(R^2_{train})$ : Median of explained variance values in the training sub-samples across folds/repetitions, $(R^2_{test})$ : Median of explained variance values in the testing sub-samples across folds/repetitions.

**Figure 4:**
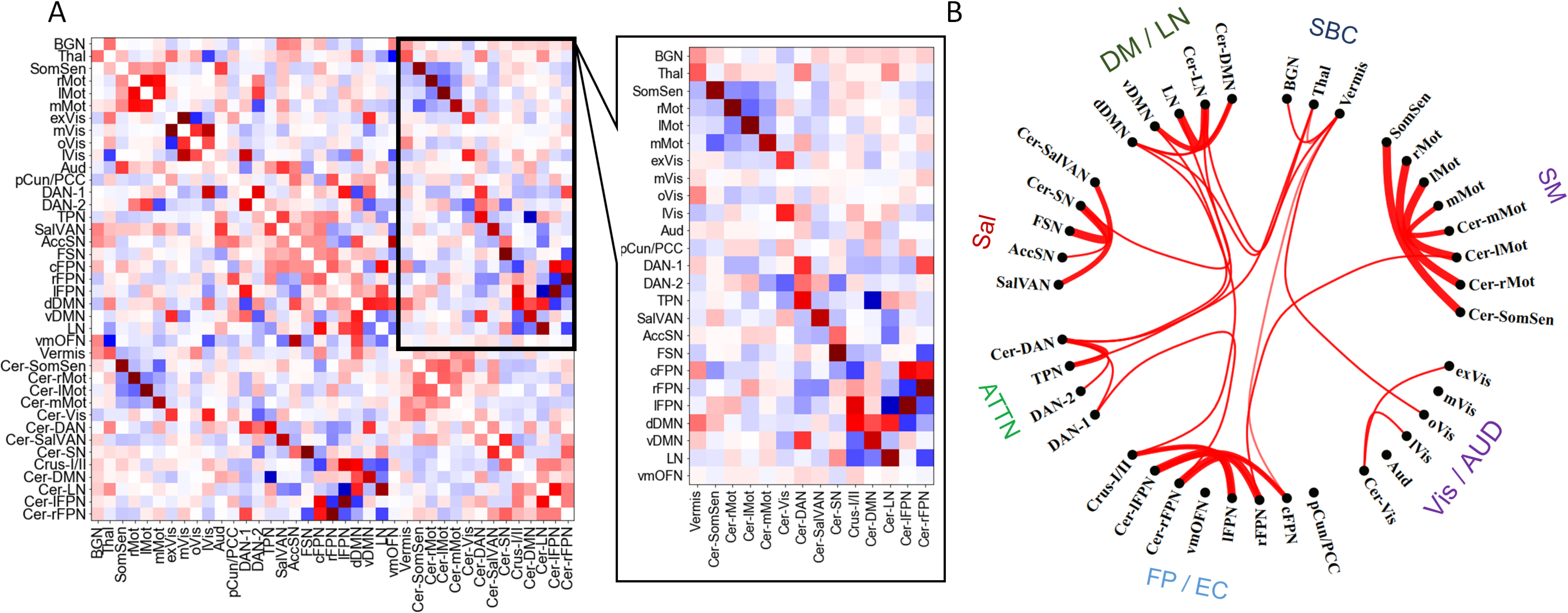
Group-average partial correlation static FC matrix and the cerebro-cerebellar submatrix. B) A circular graph showing the strongest 20% of cerebro-cerebellar edges in the group-average static FC matrix. <u>Abbreviations</u> *SBC*: Sub-Cortical networks, *SM*: Somato-motor networks, *Vis/AUD*: Visual and auditory networks, *FP/EC*: Fronto-parietal executive control networks, *Attn*: Attention networks,*Sal*: Salience networks, *DM/LN*: Default-mode and language networks

### 3.4 Cerebro-Cerebellar FC Temporal Variability and Impulsivity

The statistical details of significant associations between impulsivity and FC temporal variability are reported in table 2, whereas scatter plots are illustrated in supplementary figure S1; panels B and C. Figure S2 in supplementary material shows the inferred group-level brain FC matrices (states) along with distribution profiles of the maximum fractional occupancy values obtained from the real rs-fMRI data and the generated null data. The distribution profiles and the group average frequency of occurrence values (percentages) indicated the presence of genuine FC dynamics in the real rs-fMRI data as opposed to the null data which were mostly described by a single state.

We observed significant associations when using full correlation matrices as descriptors of dynamic brain states. Particularly, results revealed significant inverse associations between the UPPS-P lack of premeditation scale and the temporal variability of FC within the FSN-cerebellum network (*z* = −3.5, *β* = −0.34, *p<.*01, 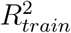 = 0.092, 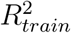 = 0.09) and the precuneus/posterior-cingulate cortex (pCun/PCC)-cerebellum network (*t* = −3.7, *β* = −0.36, *p<.*01, 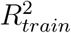 = 0.11, 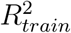 = 0.096). In addition, we identified significant associations between the UPPS-P sensation seeking scale and the temporal variability of FC within the FSN-cerebellum network (*z* = 3.3, *β* = 0.32, *p* = 0.019, 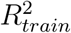 = 0.078, 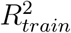 = 0.08), the pCun/PCC-cerebellum network (*z* = 3.6, *β* = 0.35, *p<.*01, 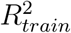 = 0.093, 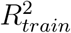 = 0.086), the BGN-cerebellum network (*z* = 3.1, *β* = 0.3, *p* = 0.037, 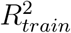 = 0.068, 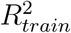 = 0.063), and the thalamus-cerebellum network (*z* = 3.3, *β* = 0.32, *p* = 0.019, 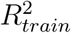 = 0.078, 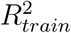 = 0.074). No significant associations were observed when using partial correlation matrices as descriptors of dynamic brain states.

## 4 Discussion

This study addressed the hypothesis that the cerebellum plays a role in impulsivity by testing possible associations between different aspects of cerebero-cerebellar rs-FC and self-reported impulsivity. We observed evidence that linked BAS, BIS, UPPS-P sensation seeking, and UPPS-P lack of premeditation to summary metrics of static and dynamic cerebro-cerebellar FC in a group of healthy young subjects. Specifically, we found that the BAS scale, which measures the tendency to engage in goal-directed and rewarding behaviors, was inversely associated with FSN-cerebellum FC strength. Likewise, we identified a significant inverse association between the BIS scale, which measured the tendency to avoid actions with negative or unpleasant outcomes, and the BGN-cerebellum FC strength. In addition, our results revealed a significant inverse association between the UPPS-P lack of premeditation, which measures the tendency to act rashly without prior reflection, and the overall FC temporal variability within the FSN-cerebellum and pCun/PCC-cerebellum networks. Finally, we found that the UPPS-P sensation seeking scale, which measures the propensity to seek novelty and experience, was associated with the temporal variability of FC within the FSN-cerebellum, pCun/PCC-cerebellum, BGN-cerebellum, and thalamus-cerebellum networks. Permutation testing and cross-validation based on repeated stratified 5-folds supported the significance and cross-validity of these findings, respectively. Taken together, the findings from this study improve current knowledge on the neural underpinnings of impulsivity, and highlight the utility of both static and dynamic FC methods in delineating the neural correlates of this and potentially other personality traits.

Impulsivity is mediated by control and reward processes, and it is likely that distinct brain networks contribute to its different forms (Whelan et al., 2012). In the context of the current findings, the implicated cerebral networks included regions known to sub-serve executive control, reward, emotion, and motivation. The frontal salience network (FSN) primarily included parts of the dorso-lateral prefrontal cortex, superior frontal cortex, fronto-polar cortex, and intra-parietal cortex (or Brodmann areas BA7, BA8, BA9, and BA10) that are known for goal-directed (top-down) selection of responses to stimuli (Corbetta and Shulman, 2002). Particularly, the dlPFC has been observed to be activated during reward anticipation and to integrate and transmit representations of reward to the mesolimbic and mesocortical dopamine systems, thus contributing to the initiation or termination of motivated behavior, whereas the fronto-polar cortex is known to be involved in decision making, reward, and attention (Ballard et al., 2011; Boschin et al., 2015). The pCun/PCC network included parts of the precuneus cortex (pCun) and dorsal sub-regions of the posterior cingulate cortex (PCC) (or Brodmann areas BA7 and BA23). The dorsal PCC is part of the large-scale fronto-parietal control network and is also connected to the default-mode and salience networks (Yeo et al., 2011; Leech and Sharp, 2013). In addition, studies have observed that the PCC is activated during tasks involving a form of monetary incentive and have indicated that it functions as an interface between motivation-related regions and top-down control of visual attention (Small et al., 2005; Engelmann et al., 2009). The basal-ganglia network (BGN) included the striatum which is regarded as a core reward region and often implicated in cue-reactivity, incentive learning, and reinforcement learning mechanisms (Elliott et al., 2000; Yin and Knowlton, 2006; Koob and Volkow, 2016). Finally, the thalamus is known for being a hub that relays sensory input towards the cerebral cortex and provide channels for the cerebellum and basal-ganglia to communicate at the cortical and sub-cortical levels (Hintzen et al., 2018). The thalamus is connected to many brain regions and is involved in response inhibition, emotion, arousal, memory, and cognitive flexibility (Taber et al., 2004; Rikhye et al., 2018; Wolff and Vann, 2019).

Recent evidence from resting-state FC studies have implicated several networks comprising key control and reward brain regions in impulsivity. One study has reported significant associations between elevated impulsivity and decreased FC between prefrontal control and sub-cortical appetitive drive regions in healthy young individuals (Davis et al., 2012). Another study has reported significant associations between basal ganglia-thalamo-cortical FC and BIS-11 motor and non-planning impulsivity in healthy adults (Korponay et al., 2017). In addition, fronto-striatal FC has been observed to be strongly associated with BAS fun-seeking (Angelides et al., 2017). Moreover, significant associations have been observed between the sensitivity to immediate or delayed reward and FC intensity within networks involving the striatum, PCC, medial and lateral prefrontal regions, and the anterior cingulate cortex (Li et al., 2013). Finally, FC between a precuneus/posterior cingulate cortex seed, the thalamus, and regions of the midbrain has been found to be associated with trait urgency (Zhao et al., 2017). These studies, among others, highlight the key role of frontal, parietal, and sub-cortical regions in the different dimensions of impulsivity.

The cerebellum is now known to be reciprocally interconnected with the aforementioned control and reward regions (Moreno-Rius and Miquel, 2017). The general role ascribed to the cerebellum is supervised learning or prediction via internal forward models (Ishikawa et al., 2016). This means that the cerebellum acts as a regulator of outcome within the networks and systems it is engaged with, in response to environmental demands and inputs (Moreno-Rius and Miquel, 2017). In this context, it has been proposed that the cerebellum may engage with prefrontal regions in restraining ongoing actions in response to external and internal stimuli, and an increased fronto-cerebellar FC strength may be necessary for improving prefrontal functionality in top-down control of goal-directed and rewarding behaviors (Miquel et al., 2019). This might explain the observed inverse association between the FSN-cerebellum FC strength, computed using partial correlations, and the BAS scale where higher values indicate a greater tendency to engage in goal-directed rewarding actions with less constraints (Gray, 1991). In addition, increased basal ganglia-cerebellar FC strength is believed to promote the initiation of motivated “Go” brain mechanisms at the expense of “No-Go” inhibitory control mechanisms which can lead to increased impulsivity (Miquel et al., 2019). This may explain the observed inverse association between BGN-cerebellum FC strength, computed using partial correlations, and the BIS scale where lower values indicate less sensitivity to unpleasant cues and outcomes. Taken together, these findings reflect a possible effect of direct cerebellar modulation of prefrontal regions and the basal-ganglia, via closed loops, on the brain’s motivational systems.

Dynamic FC studies have indicated that the FC temporal variability characterizes how different brain regions are transiently integrated and segregated across time (Calhoun et al., 2014; Lord et al., 2017). In the context of cerebro-cerebellar FC, the greater the temporal variability, the more frequent the switching of FC strength between each cerebral RSN of interest and the cerebellum. Alternatively, greater cerebro-cerebellar FC variability means that cerebellar RSNs are transiently recruited by large-scale cerebral RSNs that sub-serve complex processes. Our findings revealed an inverse association between lack of premeditation and the temporal variability of FC between the cerebellum and two top-down inhibitory control networks (FSN and pCun/PCC). In addition, sensation seeking was found to be associated with the temporal variability of FC between the cerebellum and networks that sub-serve inhibitory control, cue-reactivity, and salience attribution (BGN, thal, FSN, and pCun/PCC). These significant associations were observed only in the case of full correlation dynamic FC matrices. This indicates that the cerebellum may affect complex cognitive processes, that require dynamic interactions of multiple networks, beyond its direct closed-loop connections. This also indicates that the transient recruitment of cerebellar RSNs by these cerebral RSNs might be imperative for the modulation of complex cognitive processes that influence lack of premeditation and sensation seeking. Another possible indication might be related to the postulate that the cerebellum coordinates and links cognitive units of thought in a similar fashion to coordinating multimuscled movements (Buckner, 2013). In other words, the cerebellum may be transiently recruited to coordinate and modulate information flow within and between control and reward networks, hence indirectly influencing certain forms of impulsivity.

To our knowledge, no previous literature has thoroughly examined or reported associations between the dynamics of cerebro-cerebellar FC and behavior in healthy individuals. Therefore, interpretations of the present findings remain speculative. In addition, since only healthy subjects were included in this study, we cannot discern whether FC temporal variability within cerebrocerebellar networks is related to potentially problematic impulsive behavior. However, we speculate that cerebro-cerebellar FC variability within certain limits might be necessary for maintaining impulsive behavior in the healthy range. In addition, we hypothesize that cerebro-cerebellar FC dynamics may relate to the neural substrates of aberrant impulsive behaviors in certain brain disorders such as alcohol use disorder and attention deficit hyperactivity disorder (ADHD). Although future research is definitely needed to establish the validity of these speculations, our findings directly demonstrate, for the first time, that the cerebellum contributes to different forms of impulsivity through persistent and transient FC patterns with large-scale control and reward networks. Yet, *how* does the cerebellum dynamically regulate impulsive behavior and *how* do alterations to cerebro-cerebellar FC in certain brain disorders relate to abnormal impulsivity patterns remain important open questions.

There are several methodological limitations and concerns worth noting in the current study. A first limitation, we believe, is the use of a low-model order parcellation that only included bilateral large-scale RSNs both at the cerebral and cerebellar levels. Although this choice served our goals and interests, it was based on the fact that the HMM method is prone to over-parametrise as a function of dimensionality and may not converge reliably when the number of considered regions becomes larger (Karapanagiotidis et al., 2018; Vidaurre et al., 2017). Developing new methods that can reliably process high-dimensional data is needed in order to overcome this limitation. A second limitation is the use of positively weighted edges to estimate FC strength and temporal variability, while discarding negatively weighted edges. Negative correlations are often not reported or discarded in most studies due to the ambiguity and controversy surrounding their nature and means of analysis (Hallquist and Hillary, 2018). In addition, it is still unknown what negative correlations mean in terms of cerebro-cerebellar FC. More investigation into the nature of negative correlations and their impact on cerebellar functioning is needed in future research. A third limitation of this study is the lack of objective measurements of impulsivity such as tasks, which could have provided a better perspective with less bias than self-reported measures. Finally, future investigation into the role of the cerebellum in impulsivity and other related traits should explore gender and age differences and include clinical groups that exhibit impulsive symptomatology such as alcohol dependent individuals. This can provide further insight into the cerebellar contributions to dissociated brain functions that are altered in impulsive disorders, and that might possibly inform treatment procedures.

## 5 Conclusion

The hypothesis of a role of the cerebellum in impulsivity was investigated using a resting-state fMRI data, self-reports of impulsive behavior, and robust techniques from static and dynamic FC analysis. We have provided new evidence of a possible contribution of the cerebellum to the behavioral inhibition/approach systems, lack of premeditation, and sensation seeking. By considering how measures of cortico-cerebellar and sub-cortico-cerebellar FC relate to impulsivity in healthy individuals, this work can be extended to include clinical groups exhibiting abnormal impulsivity patterns such as substance dependence and ADHD. Taken together, our findings improve current knowledge on the neural substrates underlying impulsivity in humans and might provide a starting point for future studies. Finally, we believe that future research should include multiple groups, more variables, and task paradigms that could provide a more holistic view of the neurobiology of impulsivity and other related personality traits.

## 6 Acknowledgments

The authors thank professor Joel Swendsen for valuable advice and proof reading of the manuscript. This work was supported by a research grant fully funded by the translational research and advanced imaging laboratory (TRAIL), University of Bordeaux, France.

