## Supplementary Material for "Static and Dynamic Aspects of Cerebro-Cerebellar Functional Connectivity are Associated with Self-reported Measures of Impulsivity: A Resting-State fMRI Study"

### 1 Behavioral Variables

**Table 1:** Partial correlations between Impulsivity Variables

| | PosUrg | NegUrg | PreMed | Persev | SenSeek | BIS | BAS |
|---|---|---|---|---|---|---|---|
| PosUrg | 1 | 0.68*** | 0.16 | 0.16 | 0.00 | -0.03 | 0.10 |
| NegUrg | | 1 | 0.05 | 0.08 | -0.03 | 0.23** | -0.02 |
| PreMed | | | 1 | 0.28*** | 0.23** | -0.22** | 0.00 |
| Persev | | | | 1 | 0.04 | 0.16 | -0.34*** |
| SenSeek | | | | | 1 | -0.13 | 0.26*** |
| BIS | | | | | | 1 | -0.00 |
| BAS | | | | | | | 1 |

Abbreviations *PosUrg*: Positive Urgency, *NegUrg*: Negative Urgency, *Premed*: Lack of Premeditation, *Persev*: Lack of Perseverance, *SenSeek*: Sensation Seeking, *BIS*:Behavioral Inhibition System, *BAS*: Behavioral Approach System.

$^{*}p < .05$ , $^{**}p < .01$ ,$^{***}p < .001$

### 2 Scatter plots of significant findings

Supporting figure 1 illustrates the scatter plots that represent significant associations between different aspects of cerebro-cerebellar FC and different measures of impulsivity.

### 3 Group-level dynamic FC states

### 4 Hidden Markov model stability analysis

Previous studies that have used hidden Markov modelling of brain dynamics roughly estimated between 5 and 12 brain states (Baker et al., 2014; Ou et al., 2015; Vidaurre et al., 2017, 2018; Kottaram et al., 2019; Karapanagiotidis et al., 2018). In this study, we considered testing the stability of models with 5,6, 8, 10, and 12 states by repeating the Bayesian inference 100 times in each case

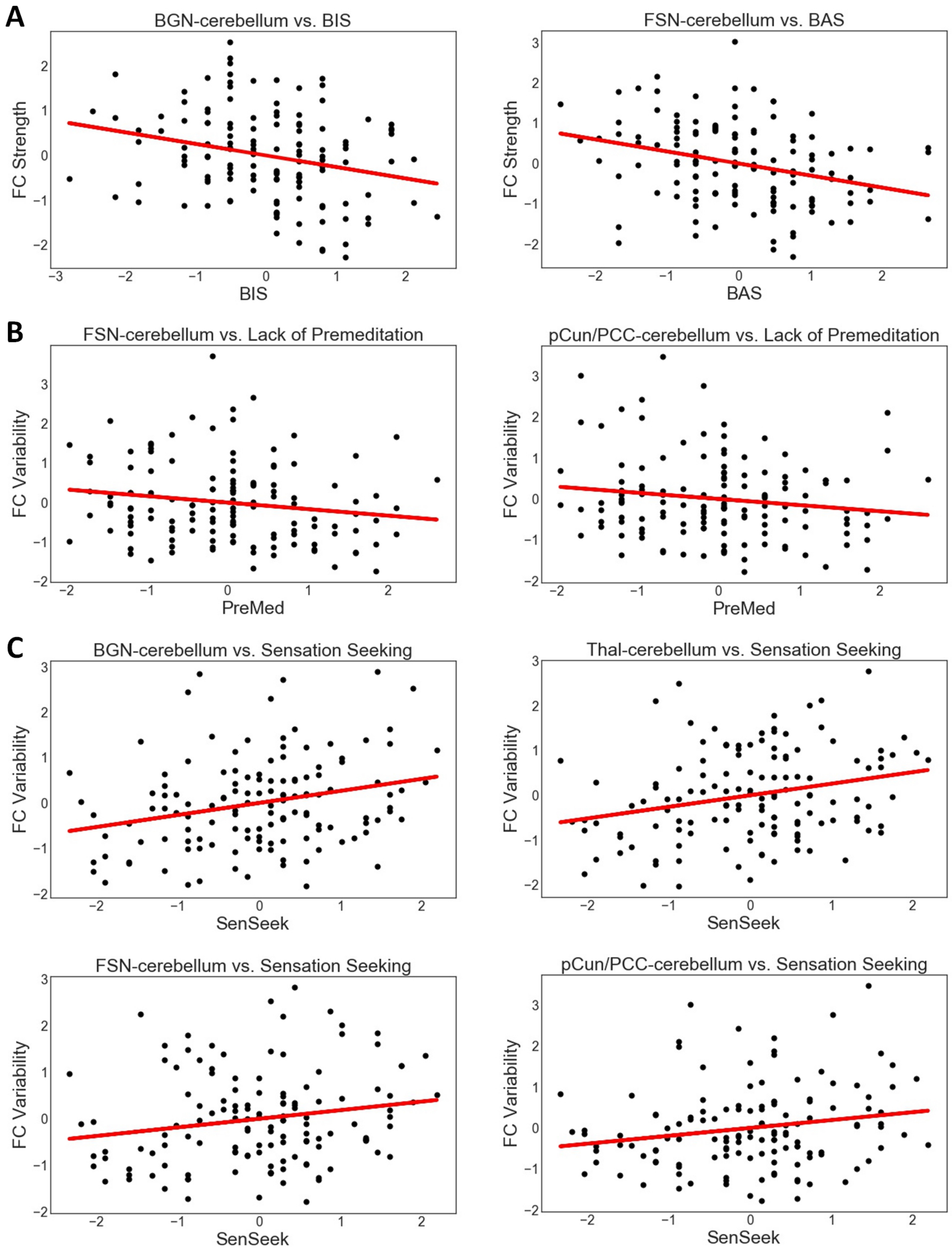


**Figure 1:** Scatter plots of significant findings. A) Cerebro-Cerebellar FC strength Vs. BIS/BAS. B) Cerebro-Cerebellar FC variability Vs. UPPS-P lack of premeditation. C) Cerebro-Cerebellar FC strength Vs. UPPS-P sensation seeking

and then estimating the degree of similarity between the states time courses across runs, measured as the amount of overlap between probabilities when the states are aligned via the Hungarian algorithm (Munkres, 1957). This was performed because the inference process can converge to different solutions

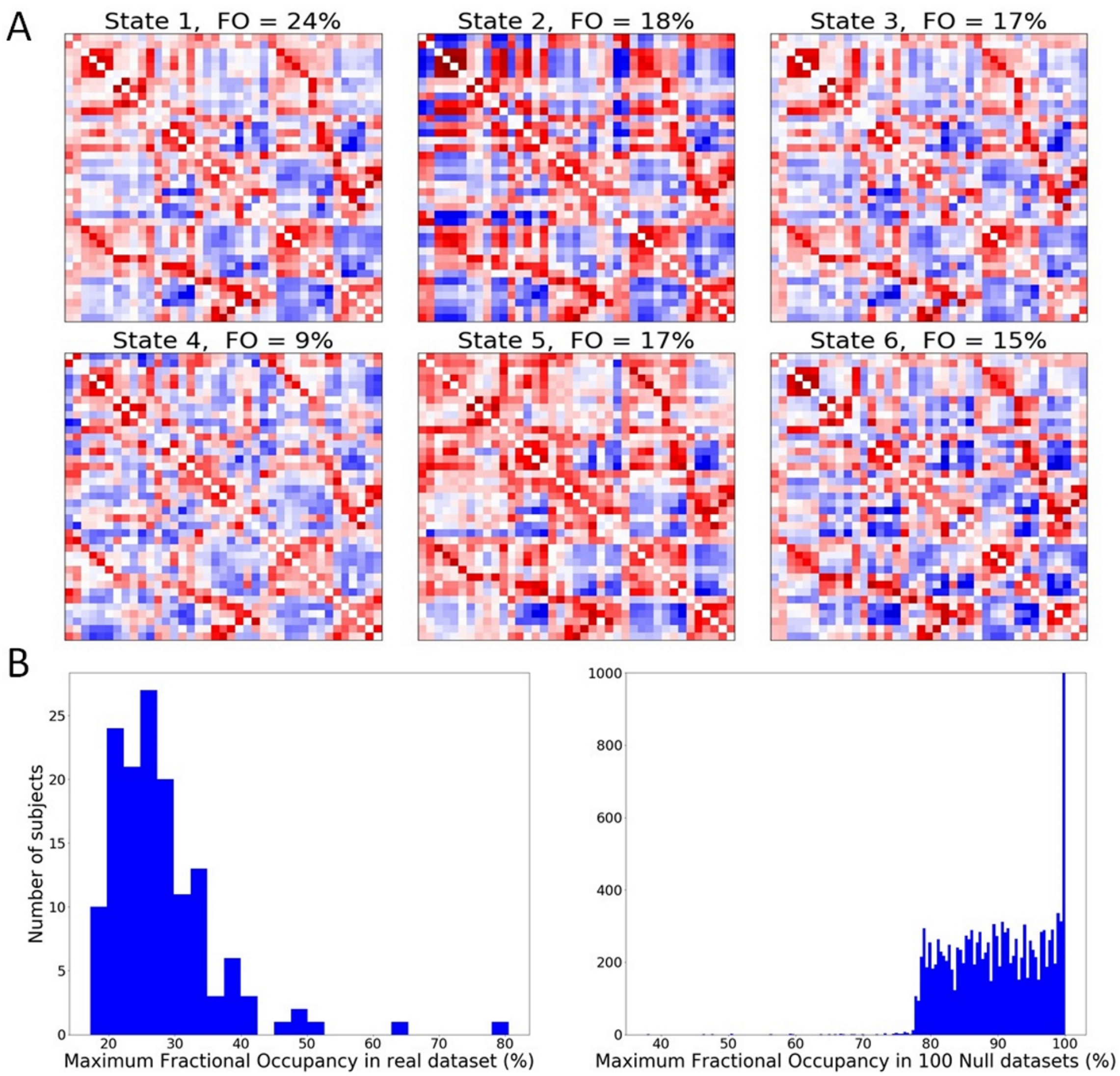


**Figure 2:** Dynamic FC analysis results. A) Group level dynamic FC patterns (states). At the top of each matrix is the group average fractional occupancy (FO) which represents the percentage of time each state is visited on average. B) Maximum fractional occupancy distributions in the real rs-fMRI dataset and the null datasets. The profiles of both distributions indicate the presence of genuine dynamics in the real rs-fMRI data.

in each run, and higher similarity between states time courses across runs indicates greater stability of the model (Karapanagiotidis et al., 2018). As expected, models with lower number of states were more stable and reproducible across runs than model with a higher number. However, high-order models attained lower levels of the free-energy index. Accordingly, we decided, as a compromise between stability across multiple runs of the algorithm, free-energy index, and temporal resolution, to use the 6 states model which was significantly more stable than the 8, 10 and 12 states models and attained lower free-energy levels than the 5 states model. Out of the 100 runs using the 6 states configuration, we chose the optimal solution that corresponded to the lowest free energy index as the final estimate of group-level dynamic brain states. Figure 2 illustrates similarity (or stability) matrices that encode levels of overlap between 100 runs of the HMM using 5 different number of states. More high values closer to 1 indicate a stable and reproducible estimation of brain dynamics across multiple runs of the algorithm. The matrices labels were re-ordered in ascending order of free-energy.

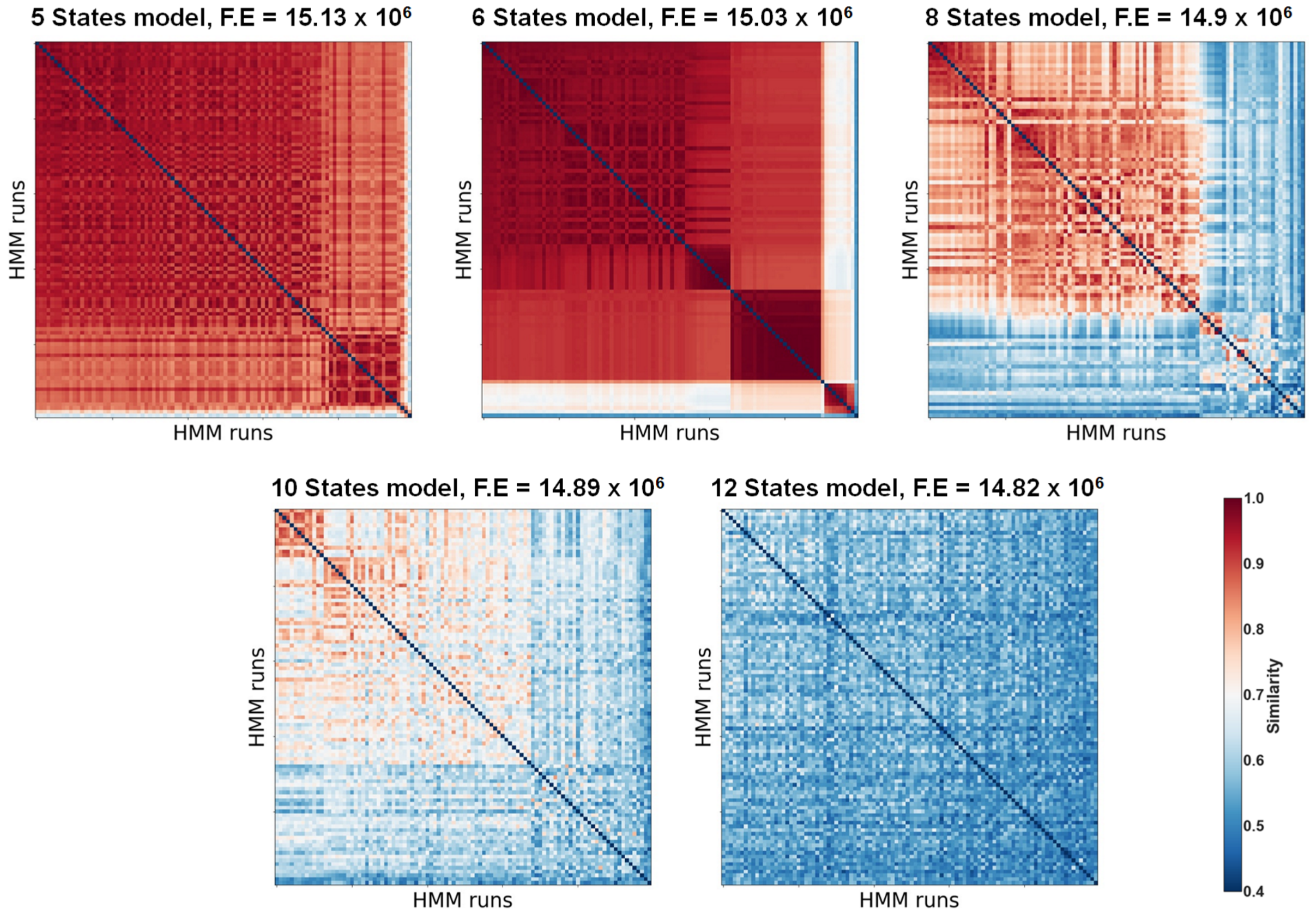


**Figure 3:** Similarity between 100 runs of HMM in 5 different configurations

### 5 Similarity between Static and Dynamic FC matrices

The rationale behind computing FC temporal variability using subject-specific dynamic FC states, was based on the observation that the static FC matrices across subjects highly resembled the frequency-weighted mean of the dynamic FC matrices: cosine similarity $> 0.98$ on average for full correlation matrices and $> 0.94$ on average when using partial correlation matrices. A bar chart showing the cosine similarity level between the static FC matrix (full correlation) and the frequency-weighted mean of the dynamic FC matrices (full correlation) at the subject-level is illustrated below in figure 3.

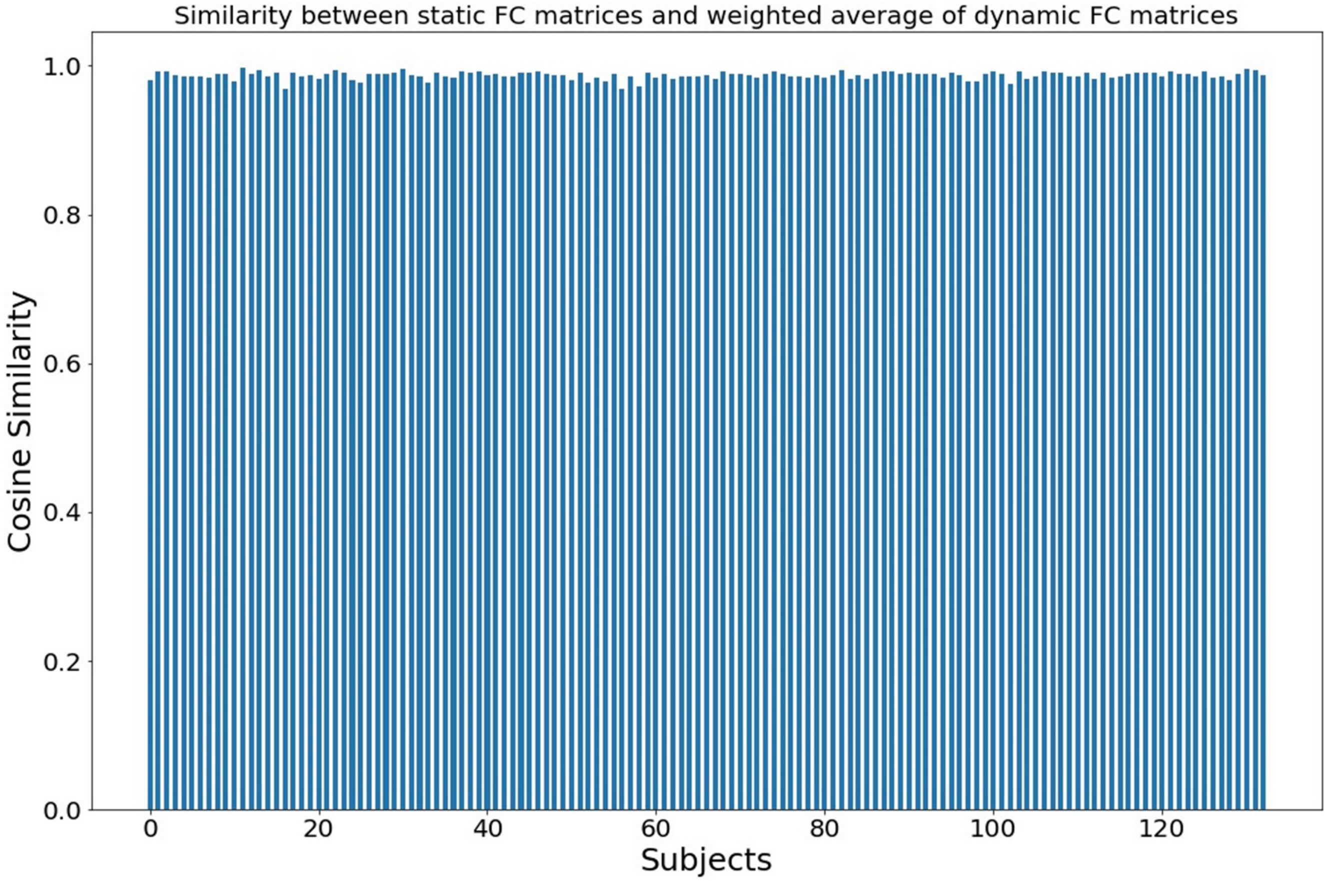


**Figure 4:** Cosine similarity between the static FC matrix and the frequency-weighted mean of the dynamic FC matrices for each subject (Mean= 0.986)

### 6 Results obtained using the 5 and 8 states HMM configurations

The results obtained using the 5 and 8 states HMM configurations reflected similar patterns of associations as those observed when using the 6 states configuration. Particularly, significant results were observed only when using full correlation matrices to describe the dynamic FC states. The statistical details of significant associations are reported in the tables 2 and 3 below. Note that we did not correct for multiple comparisons in this validation step.

**Table 2:** Results obtained using the 5 states HMM configuration

| Impulsivity scale | Network | z | $\beta$ | p-value |
|---|---|---|---|---|
| SenSeek | BGN-cerebellum | 3 | 0.29 | 0.003 |
| | Thal-cerebellum | 2.8 | 0.28 | 0.005 |
| | FSN-cerebellum | 3.1 | 0.31 | 0.002 |
| | pCun/PCC-cerebellum | 4.1 | 0.39 | <.0001 |
| PreMed | FSN-cerebellum | -3.4 | -0.33 | 0.001 |
| | pCun/PCC-cerebellum | -3.6 | -0.34 | <.001 |

**Table 3:** Results obtained using the 8 states HMM configuration

| Impulsivity scale | Network | z | $\beta$ | p-value |
|---|---|---|---|---|
| SenSeek | BGN-cerebellum | 2.6 | 0.27 | 0.008 |
| | Thal-cerebellum | 3.1 | 0.31 | 0.002 |
| | FSN-cerebellum | 2.5 | 0.26 | 0.027 |
| | pCun/PCC-cerebellum | 3.6 | 0.36 | <.001 |
| PreMed | FSN-cerebellum | -2.1 | -0.22 | 0.045 |
| | pCun/PCC-cerebellum | -2.9 | -0.28 | 0.007 |
